# THE ROLE OF ALPHA- AND BETA-ADRENERGIC RECEPTORS ON COMPULSION-LIKE ALCOHOL DRINKING

**DOI:** 10.1101/2022.11.20.517252

**Authors:** Thatiane De Oliveira Sergio, Sarah Wean, Simon Nicholas Katner, Frederic W. Hopf

## Abstract

Alcohol Use Disorders (AUD) is characterized by compulsion-like alcohol drinking (CLAD), and this intake despite negative consequences can be a major clinical obstacle. With the quite limited treatment options available for AUD, there is a significant and critical unmet need for novel therapies. The noradrenergic system is an important hub for the stress response as well as maladaptive drives for alcohol, and pre-clinical (including our own) and clinical studies have shown that drugs targeting the α1 adrenenergic receptors (ARs) may represent a pharmacological treatment for pathological drinking. However, the involvement of β ARs for treating human drinking AUD has received somewhat scant investigation, and we sought to provide pre-clinical validation for possible AR utility for CLAD. Thus, we first examined whether β AR antagonist propranolol, betaxolol (β1), and ICI, 118 551 (β2) impacted compulsion-like intake and alcohol-only drinking (AOD) in male Wistar rats through systemic injections. The systemic highest dose of propranolol (10mg/kg) reduced both AOD and CLAD. 5mg/kg propranolol affected CLAD more than AOD, with no effects of 2.5mg/kg. Similar to propranolol, betaxolol also only decreased CLAD at the lower dose (2.5mg/kg). ICI 118.551 had no effects, suggesting propranolol regulates alcohol intake through β1. Also, while AR compounds might have utility for AUD, these compounds can also lead to undesirable cardiovascular system side effects; thus, any strategy incorporating lower doses of these compounds to reduce drinking could have broad utility. Importantly, here we found that a combination of ineffective doses of propranolol and prazosin administrated together did reduce both CLAD and AOD. Finally, we investigated the effect of propranolol and betaxolol into two brain areas related to pathological drinking, the anterior insula (aINS) and medial prefrontal cortex (mPFC). Surprisingly, propranolol (1-10μg) in aINS or mPFC did not affect CLAD or AOD (although with a trend for aINS betaxolol to impact CLAD), suggesting propranolol regulation of alcohol drinking through a target other than aINS and mPFC. Together, our findings provide new pharmacological insights into noradrenergic regulation of alcohol consumption, which may inform AUD therapy.

## 1. INTRODUCTION

Alcohol Use Disorder (AUD) ranks among the most prevalent mental disorders and is characterized by compulsive heavy alcohol use and loss of control over alcohol intake, with negative consequences on both physical and mental health (Larimer et al., 1999; Koob et al., 2010; Hopf et al., 2014; Carvalho et al., 2019; G.B.D. Collaborators, 2020; Epstein, 2020). Also, the compulsion for alcohol, where consumption persists besides negative consequences, can often relate to negative affect and activation of brain stress systems (Koob et al., 2014). In addition, currently approved medications for AUD show modest therapeutic efficacy, and there is a critical need for novel therapies, especially drugs that are already FDA approved and could be quickly repurposed (Spanagel, 2009; Kranzler et al., 2018; Downs et al., 2022). However, even with robust evidence for an important relation between AUD and stress activation, there are no approved medication targeting the brain stress system to modulate excessive alcohol drinking (Haass-Koffler et al., 2018; Downs et al., 2022; Varodayan et al., 2022).

The noradrenergic system is an important hub for the stress response system (Valentino et al., 2008; Downs et al., 2022, see Discussion), as well as maladaptive drives for alcohol. Preclinical and clinical studies have shown that functional inhibition of α1 adrenergic receptors (AR) may represent a therapeutic target for AUD (Simpson et al., 2018; Sinha et al., 2021; Sinha et al., 2022). In humans, two recent trials found that the α1 AR antagonist prazosin reduces drinking and craving in patients with more severe alcohol withdrawal symptoms (Sinha et al., 2021), and alcohol withdrawal effects on the prefrontal-striatal area are also reduced by prazosin (Sinha et al., 2022). In parallel, using an animal model for compulsion-like alcohol drinking (CLAD), with alcohol adulterated with quinine, our recent findings show that systemic administration of prazosin reduces both alcohol-only drinking (AOD) and CLAD in male rats (De Oliveira Sergio et al., 2021). We also examined the impact of prazosin in the anterior insula (aINS), a key potent regulator of emotional states and a strong contributor to many aspects of addiction and emotion in humans and rodents (Centanni et al., 2021; Sommer et al., 2022). Interestingly, prazosin injection into anterior insula (aINS) also affects both CLAD and AOD, while, in contrast, aINS projections to the Locus Coeruleus area mediate CLAD but not AOD (or sweet fluid intake) (De Oliveira Sergio et al., 2021). Thus, while α1 AR modulation can influence both CLAD and AOD, some aspects of noradrenergic signaling may be more selective for challenge-related action. Similarly, we previously found that CLAD but not AOD requires aINS inputs to striatum, and also striatal inputs from mPFC (another area that can regulate drinking, see Discussion), and compulsion-like responding for alcohol activates a similar insular circuit in heavy drinking humans (Grodin et al., 2018) (see also Arcurio et al., 2015), validating the importance of these circuits for at least some aspects of problem drinking in humans.

The involvement of β ARs for AUD is least investigated (Haass-Koffler et al., 2018; Downs et al., 2022). Early clinical studies found that non-selective β AR antagonists propranolol and atenolol can reduce symptoms related to alcohol withdrawal and cravings (Carlsson et al., 1971; Zilm et al., 1975; Horwitz et al., 1989; Bailly et al., 1992). In rodents, propranolol but not nadolol reduces alcohol consumption in alcohol-dependent rats, suggesting that central, but not peripheral, β ARs are involved in regulating alcohol consumption (Gilpin et al., 2010). Furthermore, a higher dose of propranolol decreases alcohol intake in dependent P rats during early withdrawal (Rasmussen et al., 2014).

Given that compulsion is an important feature of AUD, the relation of CLAD with stress, and our previous work showing α1 AR importance for CLAD and AOD, here we hypothesized that β AR signaling would also likely be involved in regulating CLAD and AOD. We first investigated if systemic administration of propranolol (2.5, 5 and 10mg/kg), the β1 AR antagonist betaxolol (2.5 and 5mg/kg), or the β2 ARs antagonist ICI 118, 551 (1 mg/kg), would alter alcohol intake. Interestingly, we found that the intermediate dose of propranolol (5mg/kg) and lower dose of betaxolol tested (2.5mg/kg) primarily only impacted CLAD, while the highest dose of propranolol (10mg/kg) reduced both CLAD and AOD. However, the lower tested dose of propranolol (2.5mg/kg) and ICI 118,551 (1mg/kg) had no effect on drinking. Further, while clinical and preclinical studies show the beneficial effects of α1 and β antagonists on AUD, these compounds can also lead to undesirable side effects on blood pressure and cardiovascular system (Vazey et al., 2018; Downs et al., 2022). Thus, any strategy lowering the dose of these compounds to reduce drinking could have broad utility. Importantly, we found that co-administering ineffective doses of propranolol and prazosin reduced CLAD and AOD. Finally, we also investigate the role of β AR signaling in aINS and mPFC, two brain areas related to AUD and CLAD, through the direct injection of propranolol and betaxolol. Together our results provide pharmacological and neurocircuitry insights into ongoing AUD clinical trials for a novel treatment strategy that already is FDA-approved and could affect alcohol consumption.

## 2. MATERIALS AND METHODS

All experimental procedures were conducted in accordance with the Guide for Care and Use of Laboratory animals provided by the National Institutes of Health and approved by the Institutional Animal Care and Use Committee of Indiana University. All efforts were undertaken to reduce the number of animals needed and to minimize pain and suffering.

### 2.1 Subjects and Alcohol Drinking Methods

Male Wistar rats (Envigo) 45-50 days old were singly housed with *ad libitum* food and water. After two weeks of acclimatization to the vivarium, rats had access to alcohol (20% v/v diluted in water) in the intermittent two-bottle choice paradigm (IA2BC) (with the second bottle containing water). Briefly, three times a week (starting Monday, Wednesday and Friday), rats had an 18-24hr period where alcohol was available concurrently with water. The alcohol and water bottle positions were alternated across days to prevent a position bias. Intermittent access continued for ~12 weeks, as longer-term intake is necessary to facilitate development of aversion-resistant alcohol intake (Hopf et al., 2010; Seif et al., 2013; Seif et al., 2015; Spoelder et al., 2015; Spoelder et al., 2017). After ~3 months of IA2BC, rats were then shifted to limited daily access two-bottle choice (LDA), with 20 min access to 20% alcohol or water Monday through Friday. After at least 2-3wk LDA, rats had 2-3 alcohol-quinine sessions (with 10mg/L quinine) to habituate to the novelty of quinine in alcohol, and then were returned to AOD. These methods are as we previously published (Hopf et al., 2010; Seif et al., 2013; Seif et al., 2015; Darevsky et al., 2019; Darevsky et al., 2020; De Oliveira Sergio et al., 2021).

One week before starting the experimental sessions rats were gently handled by the experimenter, once a day, for ~5 min, to familiarize them with the experimental conditions and to reduce non-specific stress responses. Experimental days were generally carried out twice per week, with at least one day of AOD between test sessions; all other alcohol drinking days in the week involved alcohol-only.

### 2.2 Cannulas implantation and drugs infusion

After 3–4 weeks in LDA and 2-3 quinine sessions for habituation, surgery was performed to bilaterally implant guide cannulae (Plastics One; 26 gauge) aimed 1mm above the aINS (AP +2.8, ML ± 4.8, and DV −4.7 mm) or mPFC (AP +3.2, ML ± 1.2, and DV −3.0 mm with a 10° angle) (as in Seif et al., 2013; Seif et al., 2015; De Oliveira Sergio et al., 2021). After 7 days of recovery rats returned to LDA and had 2-3 quinine sessions again. Animals were then randomized to receive drug or vehicle during experimental sessions. As described in (De Oliveira Sergio et al., 2021), during a test session the injection needle was connected to a 10μl microsyringe (Hamilton 701-RN, USA) through a polyethylene tube. The injection needle (Plastics One; 33 gauge) was lowered to reach 1 mm below the lower end of the cannulae. To control the volume and duration of injections, a digital syringe pump (KD Scientific Inc., USA) was programmed to inject a volume of 0.6 μl at 0.2 μl/min of drugs. To reduce reflux, needle was left in place for 1 min before removal.

### 2.3 Reagents

Propranolol hydrochloride was obtained from Tocris. Prazosin hydrochloride, Betaxolol hydrochloride and ICI 118,551 hydrochloride were all from Sigma-Aldrich (USA). All drugs were dissolved in sterile saline (0.9%) except prazosin that was dissolved in sterile water. The references for the doses chosen for each compound are described in Supplemental Table 1. All drugs were prepared at the same day minutes before beginning of experiments.

### 2.4 Overview of experimental design during drinking sessions

For each study, experimental conditions were randomized across animals and sessions in a Latin-square design. For all systemic administrations (i.p), rats were habituated to the experimenter for 1-3 sessions of habituation to vehicle injection prior to injections. In study 1, propranolol or vehicle were injected 20 minutes before drinking test sessions at 2.5mg/kg (*n*=10; **Fig.1A**), 5mg/kg (*n*=16; **Fig.1B**) or 10mg/kg (n=10; **Fig.1C**). Some animals were tested with the three doses. Also, only for the propranolol systemic administration, rats were exposed twice to each condition, all conditions (e.g. vehicle vs drug) were randomized for one round, then all conditions were randomized again for a second round. For each experimental condition from a given animal, drinking data from the two rounds were averaged to give a single intake value. We routinely utilize this approach to reduce variability in our drinking measures (Seif et al., 2013; Seif et al., 2015; De Oliveira Sergio et al., 2021).

**Figure 1:**
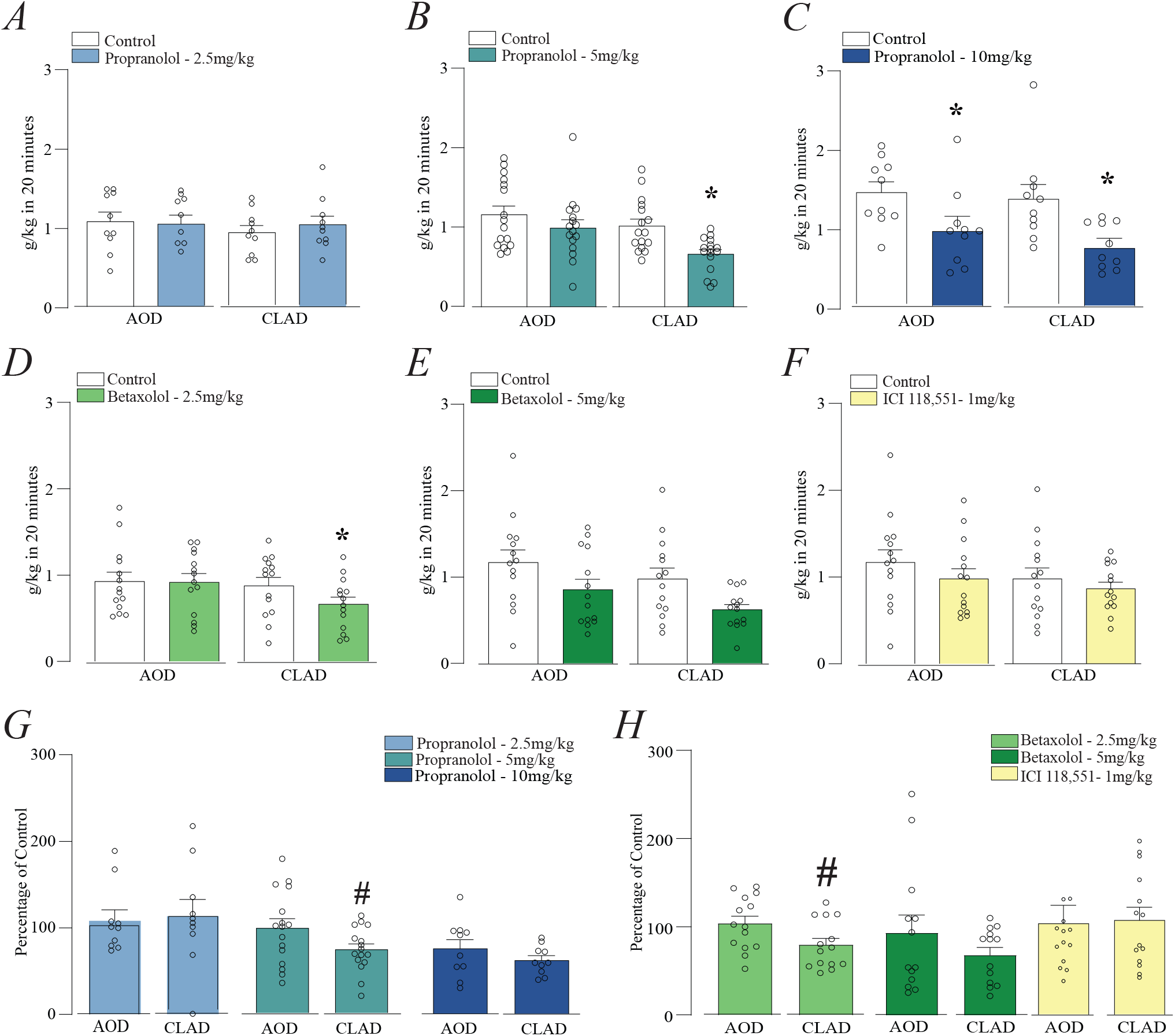
Dose dependent effect of inhibition β ARs and β1 AR with propranolol and betaxolol affects AOD or CLAD. **(A)** Systemic administration of propranolol at 2.5mg/kg did not significantly reduced AOD or CLAD. **(B)** Administration of propranolol at 5mg/kg significantly reduced CLAD but not AOD. **(C)** Administration of propranolol at 10mg/kg reduced both AOD and CLAD**. (D)** Systemic administration of β1 AR antagonist betaxolol at 2.5mg/kg reduced CLAD but did not affect AOD. **(E)** Inhibition of β1 ARs with 5mg/kg of betaxolol did not affect AOD or CLAD. **(F)** Systemic administration of β2 ARs antagonist ICI 118,551 at 1mg/kg did not affect AOD or CLAD. **(G)** Percentage of vehicle change of propranolol at 2.5, 5 and 10mg/kg showed that at the intermediated dose systemic administration reduced CLAD but not AOD. **(H)** Percentage of vehicle change for betaxolol 2.5, 5mg/kg and ICI 118,551 showed that 2.5mg/kg of betaxolol reduced CLAD but not AOD. * p<0.05; #p p<0.05

Another group of animals were injected with BTX or vehicle at 2.5mg/kg (n=14; **Fig.1D**), 5mg/kg (n=15; **Fig.1E**) and ICI 118,551 or vehicle 1mg/kg (n=15; **Fig.1F**) 30 minutes before the drinking sessions. For co-administration of an ineffective dose of prazosin (0.25mg/kg) and an ineffective dose of propranolol (2.5mg/kg) rats were first injected with prazosin or vehicle (water), and 10 minutes later to propranolol or vehicle (saline). 20 minutes after the last injection, rats went to drinking session (n=13; **Fig.3B**).

**Figure 2:**
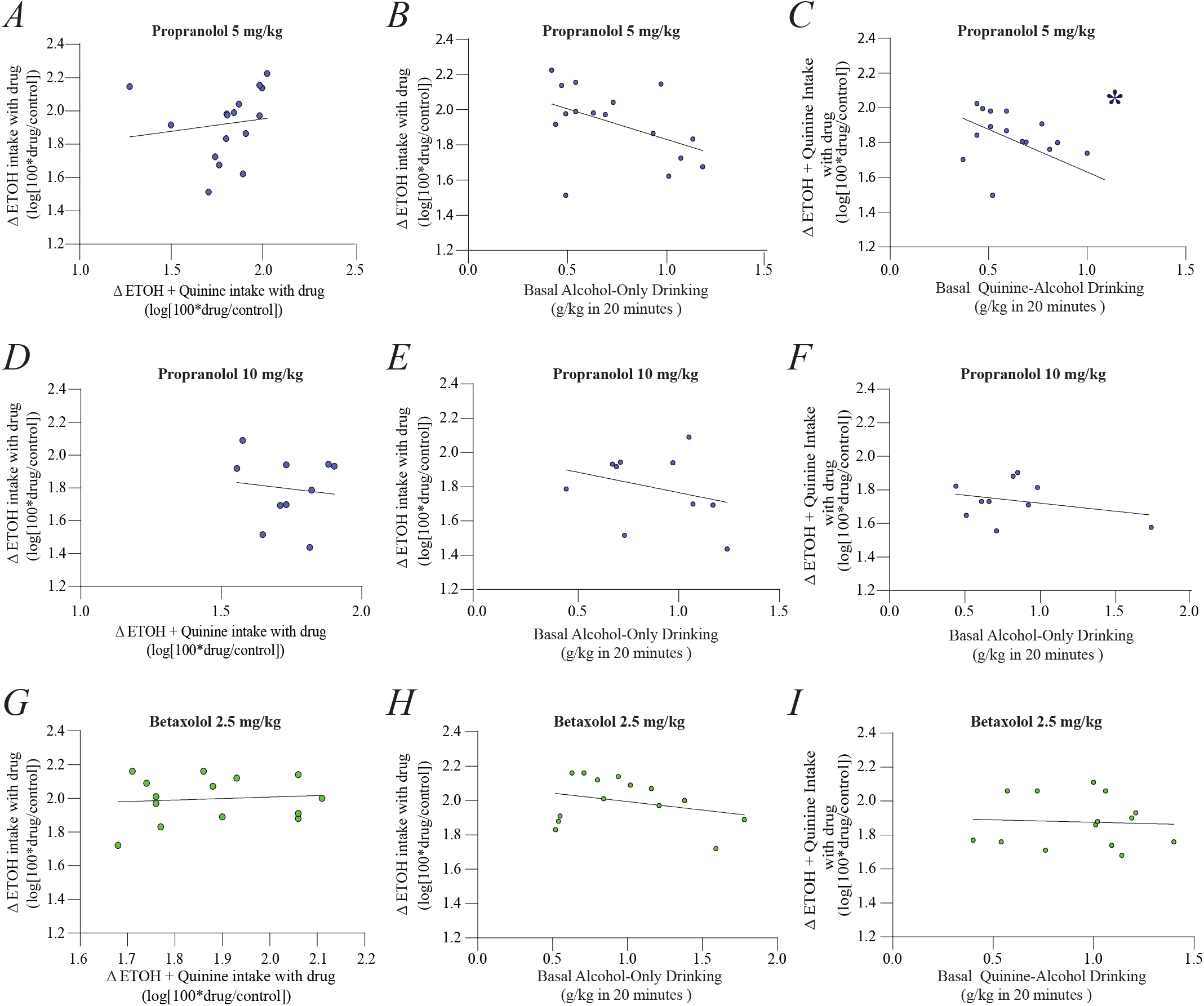
Correlations among systemic propranolol and betaxolol effects on drinking. Neither change of propranolol 5mg/kg on AOD and CLAD **(A)** or AOD basal intake **(B)** were correlated. **(C)** Changes of propranolol with basal AOD intake were negatively correlated after exclusion of one outlier (see results session and Suppl. Materials for figure with outlier). Neither changes on CLAD and AOD **(D)** basal AOD **(E)** nor basal CLAD **(F)** were correlated for propranolol 10mg/kg. Neither changes in CLAD or AOD **(G)** nor basal AOD **(H)** or basal CLAD **(I)** were correlated for betaxolol 2.5mg/kg. * p<0.05

**Figure 3.**
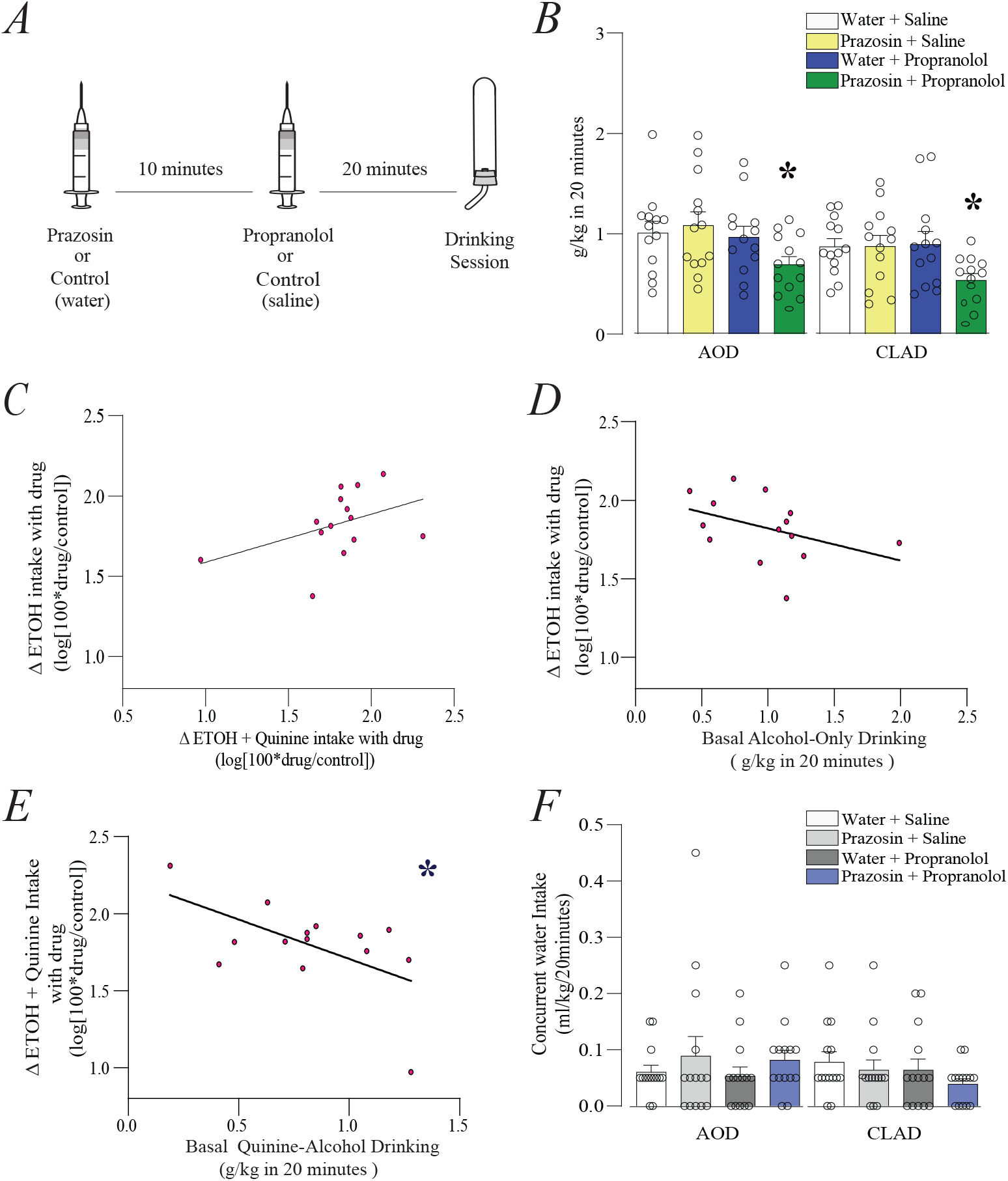
Co-administration of ineffective doses of prazosin and propranolol reduced AOD and CLAD. **(A)** Schematic experimental timeline for co-administration of ineffective doses of prazosin and propranolol. Prazosin (0.25mg/kg) or vehicle was first injected, and 10 minutes animals were injected with propranolol (2.5mg/kg) or vehicle and then 20 minutes later were exposed to AOD or CLAD drinking. **(B)** Systemic administration of prazosin or propranolol alone did not affect AOD or CLAD, however the co-administration of these drugs together decreased both drinking conditions. **(C)** Changes on AOD or CLAD promoted by the co-administration of prazosin and propranolol were not correlated. **(D)** Changes on AOD were not correlated with the basal AOD. **(E)** Changes on CLAD were negatively correlated with basal quinine-alcohol drinking. **(F)** Co-administration of prazosin and propranolol did not change the concurrent water intake. * p<0.05

For bilateral intra-aINS administration, propranolol or vehicle was tested at 0.5 and 2ug/0.6ul (n=7; **Fig.4A**) or 5 and 10ug/0.6ul (n=9; **Fig.4B**) 10 minutes before drinking session, and betaxolol or vehicle at 307ng/0.6ul 30 minutes before drinking sessions (*n*=9; **Fig.4C**). For bilateral intra-m-PFC administration, propranolol or vehicle was tested at 0.5 and 5ug/0.6ul (n=7; **Fig.5A**) 10 minutes before the drinking session. To test the effect of intra-mPFC betaxolol and prazosin, rats were bilaterally injected with 307ng/0.6ul of betaxolol or vehicle (*n*=6; **Fig.5B**), or 0.3ug/0.6ul of prazosin or vehicle (*n*=7; **Fig.5C)** 30 minutes before drinking sessions.

**Figure 4.**
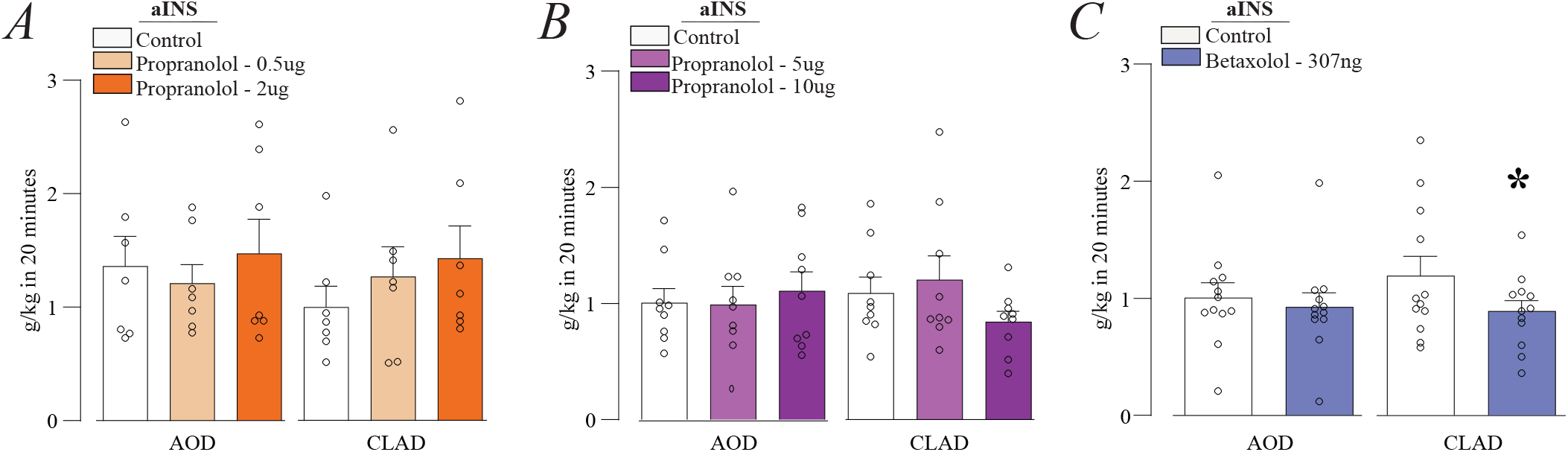
Inhibition of β ARs in aINS did not affect AOD or CLAD. Neither intra-aINS administration of propranolol 0.5, 2μg **(A)** or 5 and 10μg affect AOD or CLAD intake. **(B)** the administration of β1 ARs antagonist betaxolol (307ng) into aINS had a moderate effect on CLAD (see results). * p<0.05

**Figure 5.**
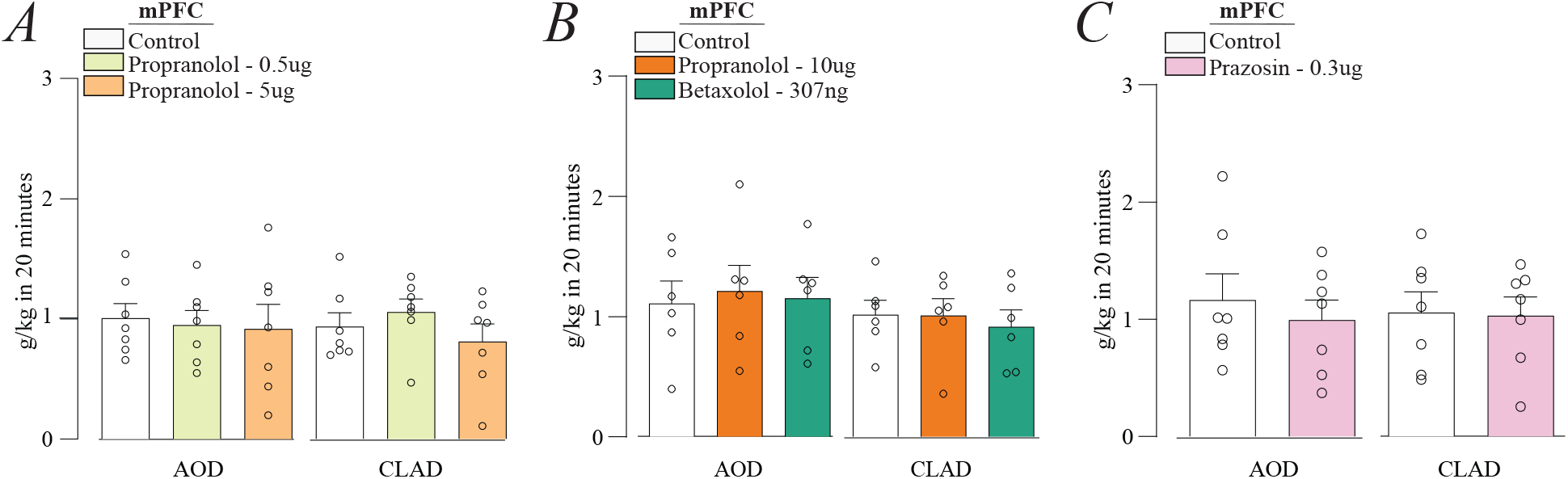
Inhibition of β and α1 ARs in aINS did not affect AOD or CLAD. Neither intra-mPFC injection of propranolol 0.5 and 5μg **(A)** or propranolol 10μg and betaxolol 307ng (B) or prazosin 0.3μg **(C)** affect AOD of CLAD.

### 2.5 Statistics and Data Analysis

Alcohol consumption was determined through changes in bottle weight before and after a drinking session and converted to grams ethanol/kilograms body weight. Concurrent water intake was determined by changes in bottle weights before and after a drinking session and expressed in ml consumption/Kilograms body weight. Statistical comparisons were primarily performed in a within-subject manner. Data were mostly analyzed by one- or two-way ANOVA with repeated-measures followed by Bonferroni test, while some comparisons used paired t-test across percentage change groups. Statistical analysis was performed using GraphPad Prism or SPSS. All data are shown as mean ± SEM.

## 3. Results

### 3.1 Effect of β_1/2_ ARs antagonists on AOD and CLAD

We first tested if systemic administration of a nonselective β AR antagonist propranolol, at the doses of 2.5, 5, and 10mg/kg, would affect CLAD and/or AOD. Our results showed that 2.5mg/kg of propranolol had no effect on CLAD or AOD [Fig.1A, n=10; two-way ANOVA; F(treatment;1,9)=0.251, p=0.628; F(drinkingcondition;1,9)=0.825, p=0.387; F(interaction;1,9)=1.046, p=0.333]. Propranolol 5mg/kg significantly decreased CLAD but had no effect on AOD [Fig.1B, n=16; two-way ANOVA; F(treatment;1,15)=11.496, p<0.004; F(drinking-condition;1,15)=19.164, p=0.001; F(interaction;1,15)=1.632, p=0.221]. Despite no interaction of drug and drinking condition with 5mg/kg of propranolol, when we normalized drinking level to vehicle for each animal, the changes in drinking were significantly greater for CLAD than AOD (Fig.1G, t_(15)_=2.455, p=0.027). However, the systemic administration of propranolol 10mg/kg decreased both AOD and CLAD, [Fig.1C, n=10; two-way ANOVA; F(treatment;1,9)=37.391, p<0.001; F(drinking-condition;1,9)=2.803, p=0.128;F(interaction;1,9)=0.262, p=0.621]. In contrast, there were no AOD versus CLAD differences in the change in drinking (drug÷vehicle) for propranolol 10mg/kg (Fig.1G, t_(9)_=1.113, *p*=0.294). Together, these data suggest that β-ARs can regulate both AOD and CLAD (evidenced by impact of propranolol 10mg/kg), while β-AR have greater regulation of CLAD (inhibited at 5mg/kg of propranolol).

Since propranolol acts through β1 and β2 ARs, we next evaluated if these dosedependent effects of propranolol on CLAD and AUD could be related to a differential activation of these two ARs. Specifically, we systemically injected betaxolol, a β1 AR antagonist or ICI 118, 551, a β2 AR antagonist, and measured AOD and CLAD intake. Our results showed that BTX 2.5mg/kg decreased CLAD but not AOD [Fig. 1E, n=14; two-way ANOVA; F(treatment;1,13)=3.578, p=0.081; F(drinking-condition;1,13)=6.341, p=0.025; F(interaction;1,13)=4.498, p=0.054].As happened with propranolol 5mg/kg, even though betaxolol 2.5mg/kg did not show interaction of drug and drinking condition althoug a trend for treatment effect, when we normalized drinking level to vehicle for each animal, the change in drinking was significantly greater for CLAD than AOD for betaxolol 2.5mg/kg (Fig.1H, t_(26)_ = 2.27, *p*=0.034). However, betaxolol 5mg/kg showed a trend to reduce CLAD but not AOD [Fig.1F, n=14; two-way ANOVA; F(treatment;1,13)=7.950, p=0.014; F(drinking-condition;1,13)=14.373, p=0.002; F(interaction;1,13)=0.057, p=0.815]. Also, there were no AOD versus CLAD differences in drug÷vehicle for betaxolol 5mg/kg (t_(13)_=0.910, p=0.370). Despite the two-way ANOVA showing a treatment effect, a t test between vehicle–AOD and betaxolol–AOD groups shows no difference (p=0.101); nonetheless, a t comparison between vehicle-CLAD and betaxolol-CLAD groups did show a significant effect (p=0.015). Thus, we assumed that there was a trend for betaxolol at 5mg/kg for affecting CLAD. Finally, 1 mg/kg of ICI 118,551 had no effect on CLAD and AOD [Fig.1D, n=14; two-way ANOVA; F(treatment;1,13)=2.557, p=0.134; F(drinking-condition;1,13)=2.591, p=0.131; F(interaction;1,13)=0.203, p=0.659], and AOD versus CLAD for drug/vehicle for ICI 118,551 was not different (t_(13)_=0.153, *p*=0.880). Taken together our findings show that the effects of propranolol on CLAD are through β2 AR and that the blocking these receptors at more middle doses reduced CLAD.

We next examined the relationship between β AR antagonist propranolol changes in AOD and CLAD across animals. We previously showed that the antagonist of α1 AR (prazosin) reduces drinking in both AOD and CLAD, but the changes in AOD with prazosin are unrelated to changes in CLAD, suggesting different α1 AR-regulated pathways for AOD and CLAD (De Oliveira Sergio et., 2021). Here we showed a related pattern for propranolol as was seen for prazosin: for 10mg/kg of propranolol, there was no relation between change in AOD and change in CLAD (Fig.2D, R^2^=0.013,*p*=0.745). There was also no correlation between 5mg/kg propranolol impact on AOD versus CLAD (Fig.2A, R^2^=0.195, *p*=0.605), although this likely reflected where the intermediate dose propranolol only affected CLAD. Further, when we examined the relationship between β1 AR antagonist betaxolol changes in AOD and CLAD across animals, we found a similar pattern, where the AOD versus CLAD change in drinking with betaxolol 2.5mg/kg showed no correlation (Fig.2D, R^2^=0.009, *p*=0.743), and no correlation for betaxolol 5mg/kg (Fig.2G, R^2^=0.019, *p*=0.605),

We also examined how basal intake might vary with drug effectiveness. For example, we find that NMDA receptor blockers (Wegner et al., 2019), prazosin (De Oliveira Sergio *et al*., 2021), and orexin receptor blockers in mice (Lei et al., 2019), are more effective in reducing alcohol drinking in individuals with higher basal intake. Here, basal intake (determined by vehicle injections day) was significantly negatively correlated with the propranolol 5mg/kg effect on CLAD (Fig.2C, R^2^=0.281, *p*=0.034). For AOD, basal intake was not correlated with propranolol 5mg/kg change in drinking for AOD (Fig.2B, R^2^=0.209, *p*=0.074), although when we removed one outlier it was significantly negatively correlated for AOD (Fig.2C, R^2^=0.504, *p*=0.030), and slopes for CLAD and AOD were close (AOD=0.129 and CLAD=0.209). However, for 10mg/kg of propranolol, there was no relation between basal intake and change in AOD (R^2^=0.088, *p*=0.404) or CLAD (R^2^=0.083, *p*=0.729). Similarly, basal intake was not correlated with BTX (2.5mg/kg) change in drinking for AOD (Fig.2E, R^2^=0.084, *p*=0.313) or for CLAD (Fig.2F, R^2^=0.004, *p*=0.813), different for what happened with propranolol 5mg/kg (Fig.2C, with bigger CLAD effect in higher drinkers). Finally, basal intake was not correlated with BTX 5mg/kg change in drinking for AOD (Fig.2H, R^2^=0.085, *p*=0.310) or CLAD (Fig.2I, R^2^=0.003, *p*=0.845). Taken together, our data suggest that impacts of different compounds have the potential to vary with basal drinking level (De Oliveira Sergio et *al*., 2021; Lei et *al*., 2019; Wegner et *al*., 2019), although we find evidence of this only for intermediate dose of (5mg/kg) propranolol.

### 3.2 Targeting α1 and β ARs to regulate CLAD and AOD

Despite our findings showing the effects of α1 and β antagonists on CLAD and AOD, these compounds can also provoke a series of undesirable side effects (see Introduction). Thus, any strategy allowing lower the dose of these compounds to reduce drinking could have broad utility. We next investigated if combining an ineffective dose of these both compounds could affect CLAD and AOD intake. The results showing that combining 0.25mg/kg of prazosin with 2.5mg/kg of propranolol decreased CLAD and AOD intake, [Fig.3B, n=14; two-way ANOVA; F(treatment;3,39)=7.753, p<0.001; F(drinking-condition;1,13)=10.722, p=0.006; F(interaction;3,39)=0.227, p=0.877]. Moreover, for prazosin+propranolol AOD changes were not correlated with CLAD changes (Fig.3C, R^2^=0.181, **p*=*=0.128). Also, basal intake was not correlated with change in drinking for AOD (Fig.3E, R^2^=0.166, *p*=0.146) however, changes in CLAD were significantly negatively correlated for basal CLAD intake (Fig.3F, R^2^=0.328, *p*=0.033). Further, the administration of prazosin + propranolol did not affect the concurrent water intake [Fig.3G, n=14; two-way ANOVA; F(treatment;3,39)=0.386, p=0.763; F(drinkingcondition;1,13)=1.018, p=0.331; F(interaction;3,39)=1.586, p= 0.208]. Thus, our data suggest that the ineffective doses of prazosin+propranolol can be used for reducing CLAD and AOD intake.

### 3.3 Impacts of inhibition of β receptor in aINS on CLAD and AOD

In order to investigate the β ARs signaling in a brain area that regulates CLAD,we next evaluated if the effects of injection into aINS of propranolol or betaxolol. We used a widely utilized doses of intracranial propranolol or BTX (see supplementary Methods). Intra-aINS propranolol (0.5 and 5 μg/side) did not affect CLAD or AOD [Fig.4A, n=7; two-way ANOVA; F(treatment;2,12)=3.574,p=0.061; F(drinking-condition;1,6)=0.565, p=0.481; F(interaction;2,12) =2.679, p=0.109]. In a separate test, propranolol (5 and 10 μg/side) did not impact CLAD or AOD [Fig.4B, n=9; two-way ANOVA; F(treatment;2,16)=1.785, p=0.200; F(drinking-condition;1,8)=0.011, p=0.917; F(interaction;2,16)=1.102, p=0.356]. However, aINS BTX (307ng/side) injection in mPFC affected CLAD but not AOD [Fig.4C, n=12; F(treatment;1,11)=11.798, p=0.006; F(drinkingcondition;1,11)=0.446, p=0.518; F(interaction;1,11) =1.922, p=0.193]. Moreover, when we normalized drinking level to vehicle for each animal that received aINS injections of BTX we found a trend for greater change in CLAD versus AOD (t_(8)_=2.229, *p*=0.074). Together, our data suggests that there was at best a moderate participation of β1 ARs in aINS signaling in CLAD regulation.

### 3.4 Impacts of inhibition of beta receptor in mPFC on CLAD and AOD

Since our data showed no effect of propranolol into aINS and a trend of betaxolol effect the involvement of β AR within mPFC (another region that regulates CLAD). Our results showed that propranolol (0.5 and 5μg/side) did not impact CLAD or AOD [Fig.5A, n=7; two-way ANOVA; F(treatment;2,12)=0.487, p=0.626; F(drinking-condition;1,6)=0.180, p=0.687; F(interaction;2,12)=0.590, p=0.569]. The higher dose of propranolol (10μg) and BTX (307ng) also did not affect CLAD or AOD [Fig.5B, n=6; twoway ANOVA; F(treatment;2,10)=0.122, p=0.887; F(drinking-condition;1,5)=12.944, p=0.016; F(interaction;2,10)=0.489, p=0.627]. Based on the lack of effects of the β ARs antagonists into mPFC, we next investigated if the prazosin, an α1 adrenergic antagonist could affect CLAD or AOD in this structure. For this we used the same dose (0.3μg/side) used in our previous results showing that prazosin decreased AOD and CLAD when administrated into aINS (De Oliveira Sergio et al., 2021). The results showed that prazosin into mPFC did not affect CLAD or AOD [Fig.5C, n=7; two-way ANOVA; F(treatment;1,6)=2.002, p=0.207; F(drinking-condition;1,6)=0.051, p=0.828; F(interaction;1,6)=0.586, p=0.473].

## 4. Discussion

Compulsion-like alcohol drinking, where consumption continues despite negative consequences, is a major obstacle to treating AUD and is related to the activation of the stress response (Koob et al., 2014; Carvalho et al., 2019). Here we investigated the effect of the β_1/2_ antagonists on CLAD and AOD in male rats through systemic injections as well into aINS and mPFC, two brain areas related to the compulsion for alcohol in animals and humans. Based on our findings with the systemic injections, we also investigated if the co-administration of ineffective doses of β ARs antagonist propranolol and α1 ARs antagonist prazosin could affect CLAD and AOD intake.

We found that there was a dose dependent effect of propranolol on CLAD and AOD intake. While 2.5mg/kg of propranolol had no effect on CLAD or AOD, the intermediate dose (5mg/kg) only decreased CLAD, and the highest dose (10mg/kg) decreased both CLAD and AOD. Interestingly, these same doses of propranolol decreased the operant alcohol reinforced responding in dependent rats with only propranolol 10mg/kg affecting the consumption in non-dependent rats (Gilpin et al., 2010). However, in alcohol dependent P rats the effects of propranolol was variable. During the early withdrawal 10mg/kg decreased ethanol consumption but 5mg/kg had no effect during early withdrawal and on alcohol intake during abstinence (Rasmussen et al., 2014). It is possible that the discrepancy of these results with our findings is related to the alcohol model used (voluntary or not) and breed of the animals (P rats versus Wistar rats). Our findings also showed that changes of the highest dose of propranolol on CLAD and AOD were not correlated, suggesting the presence of different mechanisms through β ARs regulate CLAD and AOD. Our recent study (De Oliveira Sergio et al., 2021) also showed that the systemic effects on CLAD and AOD promoted by the α1 AR antagonist prazosin were not correlated. We speculate that the noradrenergic modulation on CLAD and AOD could be regulated through different mechanisms and future studies will be necessary for better understand this process.

We next investigated if the dose dependent effect of propranolol on CLAD and AOD intake could be regulated by a differential inhibition of β1 ARs and/or β2 ARs through the injection of the antagonist of β1 ARs betaxolol and the antagonist of β2 ARs ICI 118,551. To our knowledge this is the first study to investigate the effects of betaxolol and ICI 118,551 in animals models of AUD. Our results showed that ICI 118,551 had no effect on CLAD or AOD. Moreover, betaxolol had a preferential effect on CLAD over AOD at the lowest dose (2.5mg/kg) but no effect on highest dose (5mg/kg). Taken together our findings suggests that β1 ARs signaling have a great control on CLAD and that the effect of the intermediate dose of propranolol could be related to an inhibition of β1ARs. It is known that both ICI 118, 551 and betaxolol promptly cross the blood brain barrier (Swartz, 1998; Moresco et al., 2000; Hare et al., 2006), also betaxolol is considered a highly selective antagonist for β1 ARs (Tondo et al., 1985; Satoh et al., 1993) and ICI 118,551 a selective antagonist for β2 ARs (Nathanson, 1988; Willette et al., 1999). Prior findings investigating the effects of betaxolol and ICI 118, 551 in addiction animal models have produced mixed results. The systemic administration of betaxolol at 5mg/kg during early cocaine withdrawal decreased cocaine withdrawal-related anxiety-like behavior in rats without affecting the locomotor activity (Rudoy et al., 2007). However, betaxolol had no effect on cocaine self-administration (Wee et al., 2008), and ICI 118,551 but not betaxolol (10mg/kg) blocked the stress-induced the reinstatement of cocaine in mice (Mantsch et al., 2010). Furthermore, 20mg/kg of betaxolol but not the lowest doses blocked the reinstatement of cocaine induced by stress (Vranjkovic et al., 2012). Interestingly, betaxolol and ICI alone had no effect on the compulsion-like behavior measured on nestlet shredding model, but the combination of both drugs decreased the compulsion as well as propranolol at 10mg/kg (Lustberg *et al*., 2020), suggesting β2 importance for some compulsion-related conditions.

Despite the effects of pranolol and betaxolol on CLAD intake, one significant challenge with ARs compounds is that they also regulate cardiovascular function, with potential for significant side effects (also betaxolol is FDA approved only as ophthalmic solution). Thus, the development of strategies that could decrease the doses of these drugs used for AUD could be an advantage for pharmacotherapy. We then investigated how the blocking of β ARs and α1 ARs with ineffective doses of propranolol (2.5mg/kg) and prazosin (0.25mg/kg) could affect CLAD and AOD intake. Our findings showed that the lowest doses of propranolol and prazosin alone did not affect CLAD or AOD intake, however when both drugs are combined there was a reduction on CLAD and AOD intake. Interestingly, changes on AOD and CLAD were not correlate and changes on CLAD were negatively correlated with the basal intake as happened with 5mg/kg of propranolol and showed in our previous study with 0.75mg/kg of prazosin (De Oliveira Sergio et al., 2021). Also, the concurrent consumption of water was not affected by co-administration of propranolol and prazosin suggesting no unspecific effect of the combination prazosin and propranolol on drinking. Interestingly, the ineffective dose of prazosin used here (0.25mg/kg) also did not show effect on a model of self-administration of nicotine (Forget et al., 2010). Prior results show that combining a low dose propranolol (10mg/kg) with a low dose of prazosin (1mg/kg) decreased the voluntary alcohol intake in dependent P rats during early withdrawal without affecting locomotion. It is also interesting that the effect of propranolol and prazosin together was more effective than each drug alone (Rasmussen et al., 2014). Also, co-administration of prazosin and propranolol decreases compulsion-like behavior in mice in the marble burying model, and the combination of each drug was also more effectively than each drug alone (Lustberg et al., 2020). However, our findings showed that despite the combination of prazosin and propranolol affect CLAD and AOD intake it was not more effective than either drug alone (data showed in supplemental material). It is not known how the exact mechanism through co-administration of prazosin and propranolol affects CLAD and AOD intake, but we speculate that the AOD could be mediated by α1 ARs and CLAD could be mediated by both α1 and β ARs. Future studies will need to investigate it better.

Based on the results above we investigated if the effects of propranolol and betaxolol could be me mediated by two brain areas relate to CLAD, aINS and mPFC. Our findings showed that all doses testes of propranolol into aINS (0.5, 2,5 and 10ug) did not affect CLAD or AOD, and that there was a trend for betaxolol in aINS to decrease CLAD. Furthermore, our findings showed no effect of propranolol or betaxolol into mPFC on CLAD or AUD. Similarly, 0.3ug of prazosin within mPFC did not affect CLAD or AOD intake, while a similar dose of prazosin within aINS decreases both CLAD and AOD (De Oliveira Sergio et al., 2021).

The aINS and mPFC receive noradrenergic projections from the Locus Coeruleus (Robertson et al., 2013). Although we did not find effects for propranolol injected within aINS, a prior study showed that propranolol at 2.5μg into aINS decreased the percentage of water consumption on incidental taste learning, but not the aversive taste learning (Miranda et al., 2008). Also, 5μg of propranolol injected in aINS decreased arousal induced neophobia although, the doses of 1 or 10μg had no effect (Rojas et al., 2015). The same has been shown for mPFC where prior studies also showed that propranolol in this area can control different responses. The 5μg of propranolol injected directly into mPFC affect aversive taste association as well as aversive retrieval but not during incidental taste memory formation (Reyes-Lopez et al., 2010). Also, the systemic administration of propranolol (10mg/kg) regulates the changes through mPFC that induced by stress that contribute to extinction fear impairment (Fitzgerald et al., 2015).

Although, our data found no effects of propranolol on CLAD and AOD when injected direct into aINS or mPFC, a recent study showed that propranolol injection into central amygdala decreased the alcohol intake only in dependent rats, while the prazosin injection in this area decreased the alcohol intake only in non-dependent rats (Varodayan et al., 2022). Also, the injection of propranolol into basolateral amygdala prevented reinstatement of alcohol seeking behaviors in a self-administration paradigm in rats (Chesworth et al., 2018). Thus, we speculate that the systemic effects of propranolol on CLAD and AOD intake could be mediated through amygdala regions, while the effects of prazosin, based on our recent study, could be mediated through aINS (De Oliveira Sergio et al., 2021).

This study has some limitations, first we only used male rats. Another recent study from our laboratory showed that males and females have different strategies on CLAD intake, with females being more persistent than males for high challenges of quinine intake (De Oliveira Sergio et al., biorvix) thus, it would be of value to investigate the effect of these compounds in females. However, despite these limitations the present findings provide new insights to ongoing clinical studies about the effect of the β AR antagonist propranolol on CLAD and mainly the advantage of using lowest doses of propranolol and prazosin in combination as a pharmacological strategy for AUD treatment.

## Supporting information

supplemental results

## 5. Authorship contribution

TDOS, and FWH designed the experiments, analyzed the data, and wrote a first draft of the manuscript; TDOS, SW and SK performed the experiments; all authors edited the manuscript and approved the final version.

## 6. Acknowledgements

The authors thank Sydney Hoffman, Heidi Cult, and Rebecca J. Smith for technical assistance with animal drinking and histology.

