## supplemental results for "THE ROLE OF ALPHA- AND BETA-ADRENERGIC RECEPTORS ON COMPULSION-LIKE ALCOHOL DRINKING"

**Results and Discussion of Suppl.Fig.1**:

Prior studies showed that the co-administration of prazosin and propranolol was able to decrease alcohol intake during alcohol withdrawal and after long imposed abstinence in P rats (Rasmussen et al., 2014) as well as decreased the compulsive-like behavior on marble burying test in mice (Lustberg et al., 2020). Interestingly, on both studies the effect of the co-administration of prazosin and propranolol was more effective than each compound alone. Our findings showed that the combination of ineffective doses of prazosin (0.25mg/kg) and propranolol (2.5 mg/kg) decreased AOD and CLAD. Thus, we evaluated if the co- administration of these compounds could be more effective than each compound alone. This information could be value for future clinical trials since that the combination of low doses of both compounds could reduce the chance of side effects. For this we compare the results of our previous study showing that systemic injection of prazosin at the doses of 0.75 and 1.75 mg/kg reduced AOD and CLAD (De Oliveira Sergio et al., 2021), with the current data obtained in current study with propranolol and betaxalol. The analysis showed that the percentage of control changes of prazosin + propranolol were not different for the administration of prazosin 0.75 and 1.5 mg/kg and propranolol 10 mg/kg on AOD [n 10-18 Fig. supp 1A, n = 7; one-way ANOVA; $F(treatment;3,55) = 0.288$, $p = 0.833$]or prazosin 0.75 and 1.5 mg/kg, propranolol 5 and10 mg/kg and betaxolol 2.5mg.kg on CLAD [n 10-18 Fig. supp 1B, n = 7; one-way ANOVA; $F(treatment;3,83) = 1.649$, $p = 0.156$]. Even if there were no difference on percentage of control changes for co-administration of prazosin and propranolol from the other compounds alone the data is suggesting an interesting pharmacological intervention that could reduce the side effects and consequently increase the patient adherence to the pharmacological treatment.

#### Supplemental Figures Legend

**Suppl. Figure 1.** Percentage of control change for prazosin, propranolol and betaxolol showing no difference for these compounds alone or when administrated together prazosin and propranolol. As we showed before prazosin 0.75 and 1.5 mg/kg decreased CLAD and AOD, here we express the data presented in De Oliveira Sergio et al., 2021, as percentage of control, aiming to compare the effect of these compounds alone and with the co-administration of ineffective doses of prazosin and propranolol. **(A)** There is no difference on AOD intake between prazosin at 0.75 mg/kg, 1.5 mg/kg or propranolol 10 mg/kg and the co-administration of ineffective doses of prazosin and propranolol**. (B)** There is no difference on CLAD intake between prazosin 0.75 1.5 mg/kg, propranolol 5 and 10 mg/kg, betaxolol 2.5 mg/kg and the co-administration of prazosin and propranolol.

**Suppl. Figure 2. Histology of placements**. **(A)** Example aINS placements. **(B)** Example mPFC placements. Br: bregma.

**Supp. Table 1:** Pharmacological compounds, doses used and references.

| Agent | Dose | References |
|---|---|---|
| Propranolol (i.p) | 2.5, 5 and 10mg/kg | Gilpin and Koob, 2010; Rasmussen *et al*., 2014 |
| Prazosin (i.p) | 0.25 mg/kg | Forget *et al*., 2010 |
| Betaxolol (i.p) | 2.5 and 5mg/kg | Mantsch *et al*., 2010; Rudoy and Van Bockstaele,2013 |
| ICI 118,551 (i.p) | 1mg/kg | McReynolds *et al*., 2014, Mantsch *et al*., 2010 |
| Propranolol (i.c) | 0.5, 2,5,10 ug | Funk *et al*., 1996; Rojas et al., 2015, Yamada *et al*., 2011 |
| Prazosin (i.c) | 0.3ug | Vranjkovic *et al*., 2014 |
| Betaxolol (i.c) | 307ng | Vranjkovic *et al*., 2014 |

i.p intraperitoneal
i.c intracranial

De Olivera Sergio et al., Fig. Supp. 1

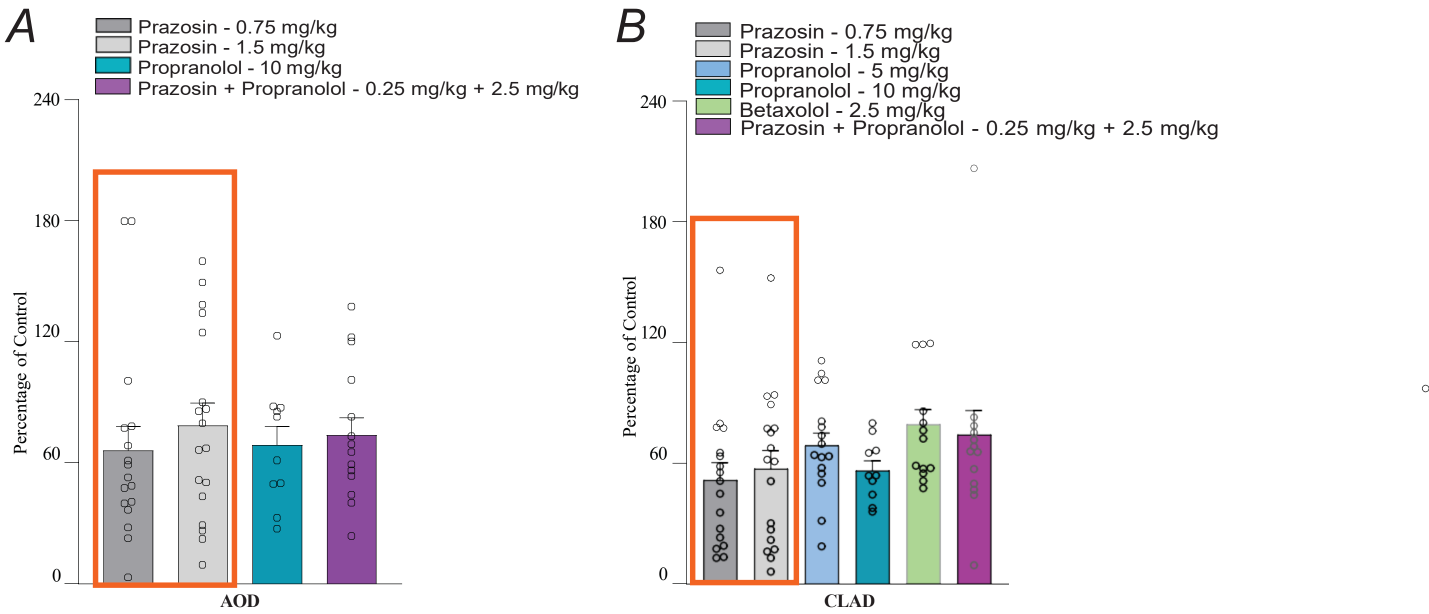

### De Oliveira Sergio et al., Fig Supp. 2

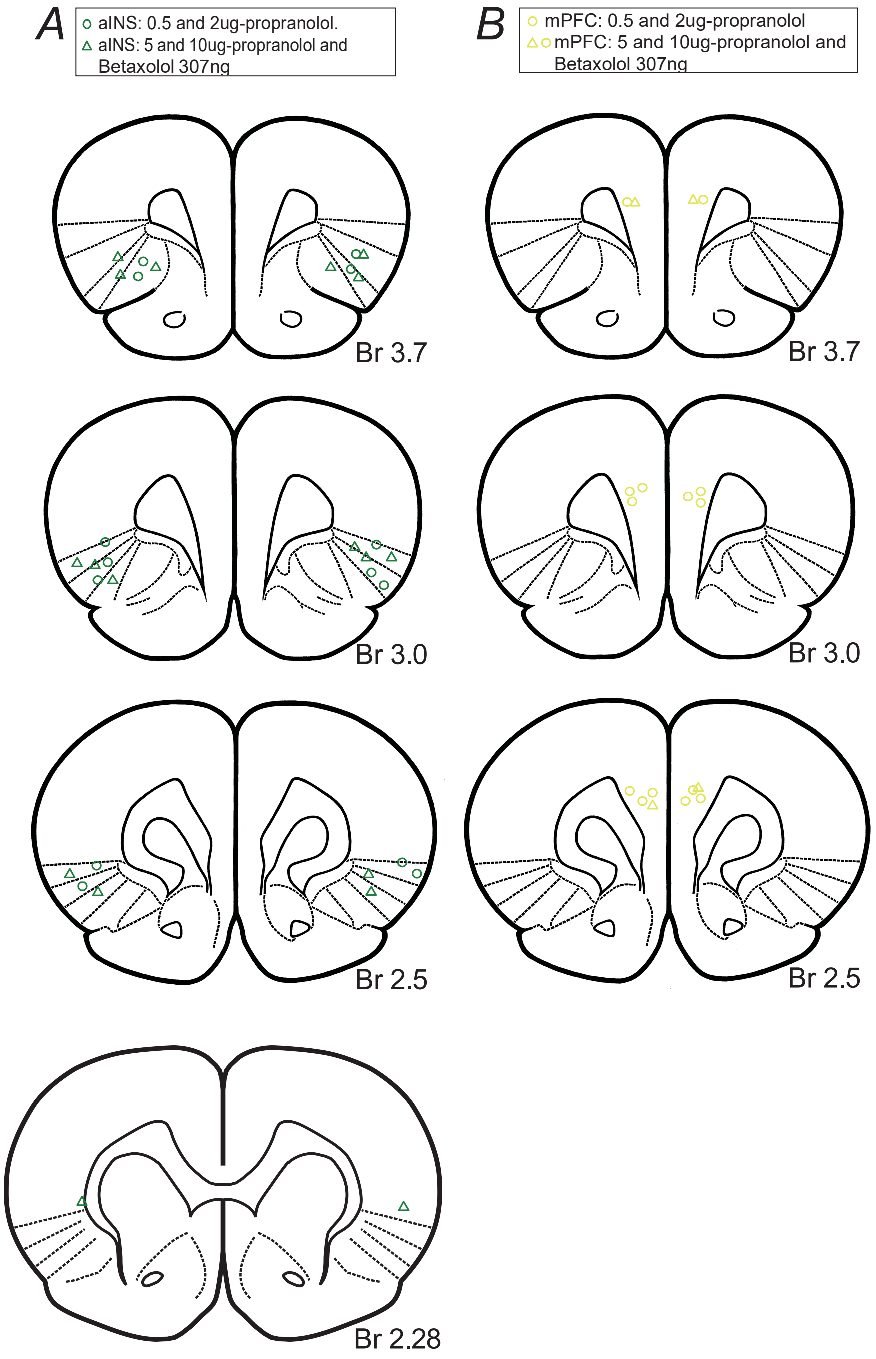
